# Microbiota-derived short chain fatty acids promote Aβ plaque deposition

**DOI:** 10.1101/2020.06.09.141879

**Authors:** Alessio Vittorio Colombo, Rebecca K. Sadler, Gemma Llovera, Vikramjeet Singh, Stefan Roth, Steffanie Heindl, Laura Sebastian Monasor, Aswin Verhoeven, Finn Peters, Samira Parhizkar, Frits Kamp, Mercedes Gomez de Agüero, Andrew J. Macpherson, Edith Winkler, Jochen Herms, Corinne Benakis, Martin Dichgans, Harald Steiner, Martin Giera, Christian Haass, Sabina Tahirovic, Arthur Liesz

**Affiliations:** German Center for Neurodegenerative Diseases (DZNE), 81377 Munich, Germany; Institute for Stroke and Dementia Research (ISD), University Hospital, LMU Munich 81377 Munich, Germany; Center for Proteomics and Metabolomics, Leiden University Medical Center (LUMC), Leiden, The Netherlands; Metabolic Biochemistry, Biomedical Center (BMC), Faculty of Medicine, Ludwig-Maximilians-Universität München, 81377 Munich, Germany; Maurice Müller Laboratories (DKF), Universitätsklinik für Viszerale Chirurgie und Medizin Inselspital, Murtenstrasse 35, Bern, Switzerland; Munich Cluster for Systems Neurology (SyNergy), Munich, Germany; Center for Neuropathology and Prion Research, Ludwig-Maximilians University Munich, 81377 Munich, Germany

## Abstract

Previous studies have identified a crucial role of the gut microbiome in modifying Alzheimer’s disease (AD) progression. However, the mechanisms of microbiome-brain interaction in AD, including the microbial mediators and their cellular targets in the brain, were so far unknown. Here, we identify microbiota-derived short chain fatty acids (SCFA) as key metabolites along the gut-brain axis in AD. Germ-free (GF) AD mice exhibit a substantially reduced Aβ plaque load and markedly reduced SCFA plasma concentrations; conversely, SCFA supplementation to GF AD mice was sufficient to increase the Aβ plaque load to levels of conventionally colonized animals. While Aβ generation was only mildly affected, we observed strong microglial activation and upregulation of ApoE upon the SCFA supplementation. Taken together, our results demonstrate that microbiota-derived SCFA are the key mediators along the gut-brain axis resulting in increased microglial activation, ApoE upregulation and Aβ deposition.

## Introduction

Alzheimer’s disease (AD) is a progressive neurodegenerative disorder characterized by the aggregation and deposition of amyloid-β (Aβ) and tau. The identification of several AD risk genes such as TREM2, CD33 or CR1 as key regulators of microglial function triggered mechanistic studies revealing the contribution of the brain’s resident innate immune cells to AD pathology. In particular, triggering microglial phagocytic clearance of Aβ can reduce amyloid plaque pathology (Hansen et al., 2018). Besides the already well acknowledged contribution of amyloidogenic protein processing and neuroinflammation to AD pathology, the gut microbiome is emerging as a novel and highly relevant modifier of brain pathology. A key function of the gut microbiome has been established over the past decade in a number of neurological diseases spanning across neurodevelopment, stroke, Parkinson’s disease and neuropsychiatric disorders (Tremlett et al., 2017). For example, we have previously described an intricate bi-directional link between the gut and brain after acute stroke, where stroke changes the gut microbiota composition (Singh et al., 2016). In turn, post-stroke dysbiosis induced changes in the immune response to stroke. Most importantly, the gut microbiota composition can be modulated to improve the disease outcome in stroke and other neurological disorders (Benakis et al., 2019), a finding that was robustly reproduced across disease entities and laboratories (Cryan and O’Mahony, 2011).

Only recently, a role of gut microbiota was also established in AD. AD patients have an altered gut microbiome compared to matched control patients, characterized by reduced species diversity and an increased abundance of Bacteroidetes (Vogt et al., 2017). Similar findings were obtained in 5xFAD and APPPS1 amyloidosis mouse models (Brandscheid et al., 2017; Harach et al., 2017). Other studies have identified that antibiotic treatment-induced changes in microbiota composition as well as microbial deficiency in GF animals was associated with reduced Aβ pathology (Dodiya et al., 2019; Harach et al., 2017; Minter et al., 2017; Minter et al., 2016). This effect could result from reduced Aβ production or increased clearance. Interestingly, GF animals as well as antibiotic-mediated dysbiotic animals exhibited reduced microglial activation (Minter et al., 2017). While previous studies documented a link between the gut microbiome and Aβ pathology, the underlying mechanisms and molecular mediators remain elusive. To address this key question, we generated a GF amyloidosis mouse model which allowed us to explore and identify bacterial metabolites that mediate gut-brain axis in AD. Moreover, we performed a detailed analysis of the gut microbiome’s impact on amyloidogenesis and neuroinflammation, in order to differentiate between direct effects on Aβ generation and effects mediated via microglial cells. Our findings identify microbiota-derived short chain fatty acids (SCFA), bacterial fermentation products of fiber, as the key mediator contributing to Aβ plaque deposition. We further identified an unexpected link between SCFA and upregulation of microglial ApoE expression that may underscore increased Aβ deposition.

## Results

### The gut microbiome promotes AD pathology

In order to study mechanisms of microbiota-brain interaction in AD, we generated a GF amyloidosis mouse model (APPPS1) by embryo transfer into axenic mice. In accordance with previous studies (Harach et al., 2017; Minter et al., 2016), we observed a striking reduction in cerebral Aβ plaque load in 5 months old GF APPPS1 animals compared to APPPS1 mice housed under conventional, specific pathogen free (SPF) conditions (**Fig. 1A**). Correspondingly, GF APPPS1 mice demonstrated a significantly better cognitive performance in a spatial memory task compared to SPF mice (**Fig. 1B**). We further analyzed the size distribution of Aβ plaques between the groups using an automated analysis paradigm of 3-dimensional plaque segmentation and quantification in the frontal cortex (Peters et al., 2018) (**Fig. 1C**). This showed that SPF mice display a significantly increased density of small (4-8 μm) Methoxy-X04-positive plaques compared to GF, while larger plaques were not affected (**Fig. 1D**). This result highlights the impact of bacterial colonization on early amyloid plaque deposition rather than plaque growth.

**Figure 1.**
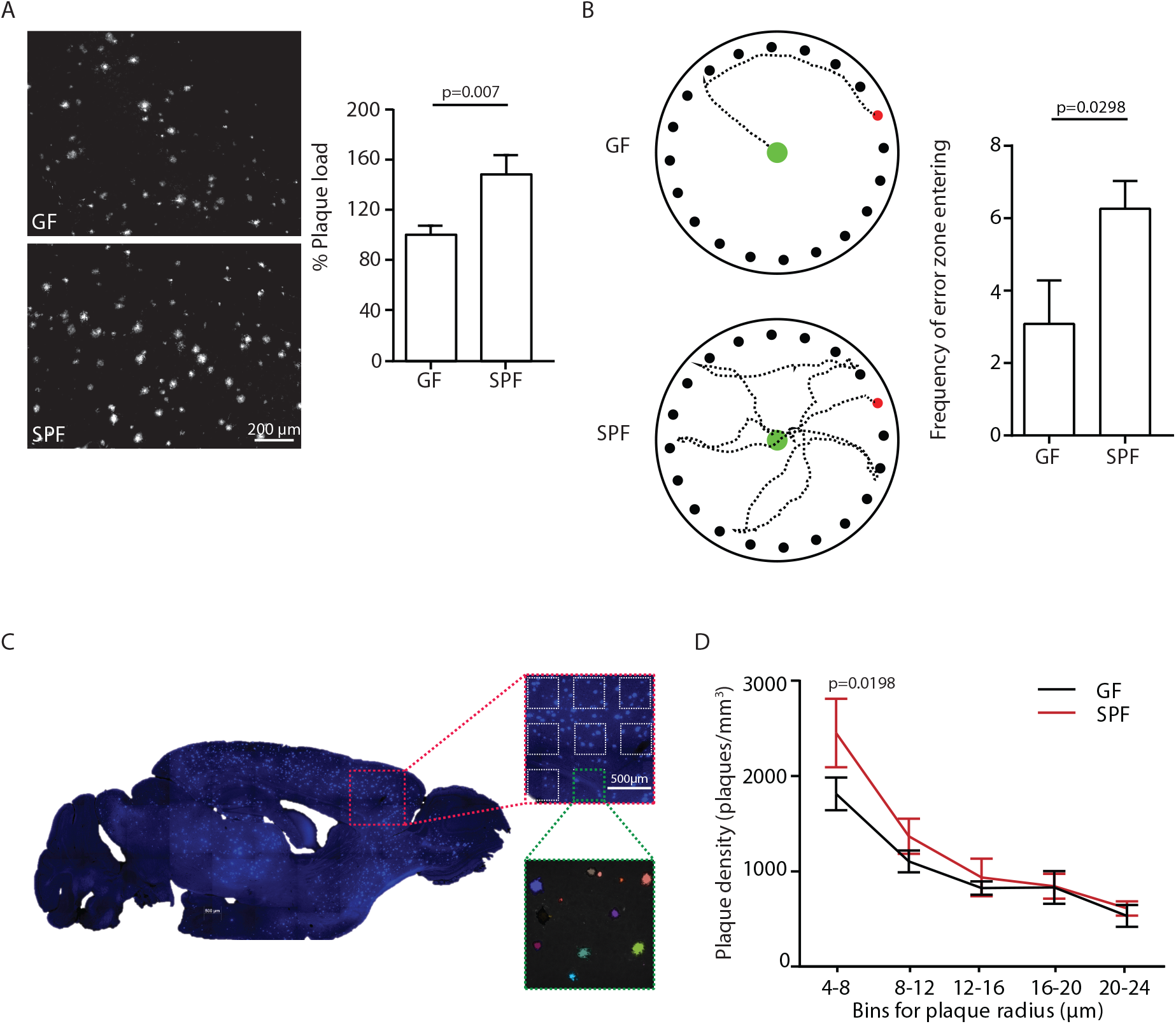
Germ-free APPPS1 mice show reduced AD pathology. **(A)** Representative images and analysis of brain cortices from 5 months old GF and SPF APPPS1 mice immunostained for Ab (clone 6E10). Quantification of parenchymal plaque load reveals significantly reduced Ab burden in GF mice compared to the SPF group. Values are expressed as percentages of amyloid plaque area and normalized to GF group (unpaired T test). **(B)** Representative results and quantification from Barnes maze behavioral analysis of 5 months old GF SPF APPPS1 mice. Quantification of frequency of error zone entering in the Barnes maze reveals a better performance in GF mice compared to SPF mice (unpaired T test). **(C)** Sagittal overview image indicating the analysis ROIs in the frontal cortex (Blue: Methoxy-X04-positive plaques) and representative image demonstrating segmentation of Methoxy-X04 fluorescence intensity into individual plaques. The images show maximal intensity projections. Individual plaques are labelled with different colors. **(D)** Frequency distribution of plaque radius in 5 months old GF and SPF APPPS1 mice (two-way ANOVA). All data presented as mean ± SEM from at least 5 mice per group.

As our initial experiments indicated a specific effect of the gut microbiome on the development of small plaques, we focused our mechanistic studies on the early phase of Aβ plaque deposition, which corresponds to the 3 months of age in APPPS1 mouse model. To identify bacterial mediators along the gut-brain axis in AD, we analyzed the blood metabolome of 3 months old GF and SPF APPPS1 animals (**Fig. 2A**). Using NMR based metabolomics, we identified 15 distinct metabolites to be regulated between GF and SPF APPPS1 mice. Among these, acetate, the most abundant SCFA, was the most regulated with a more than 7-fold higher concentration in SPF compared to GF mice. Therefore, we further quantified the total plasma SCFA concentrations of acetate, butyrate and propionate (C:2-C:4 SCFA). We detected an approx. 4-fold higher concentration of total SCFA concentrations in SPF compared to GF APPPS1 mice (**Fig. 2B**). These results suggested that SCFA may provide a pathomechanistic link between bacterial colonization and Aβ pathology. In order to study a causal link between SCFA and Aβ deposition, we performed a SCFA supplementation experiment by providing SCFA to GF APPPS1. This approach has been well-established in previous studies by us and others and demonstrated to normalize plasma SCFA levels to physiological concentrations (Erny et al., 2015; Sadler et al., 2020). Mice received SCFA in drinking water for 8 weeks, starting at 4 weeks of age (**Fig. 2C**). Strikingly, SCFA supplementation of GF APPPS1 mice was sufficient to nearly double cerebral Aβ plaque load (**Fig. 2D**) Taken together, our results demonstrate that SCFA are the key effector molecules that are sufficient to mediate effects of bacterial gut colonization onto Aβ pathology.

**Figure 2.**
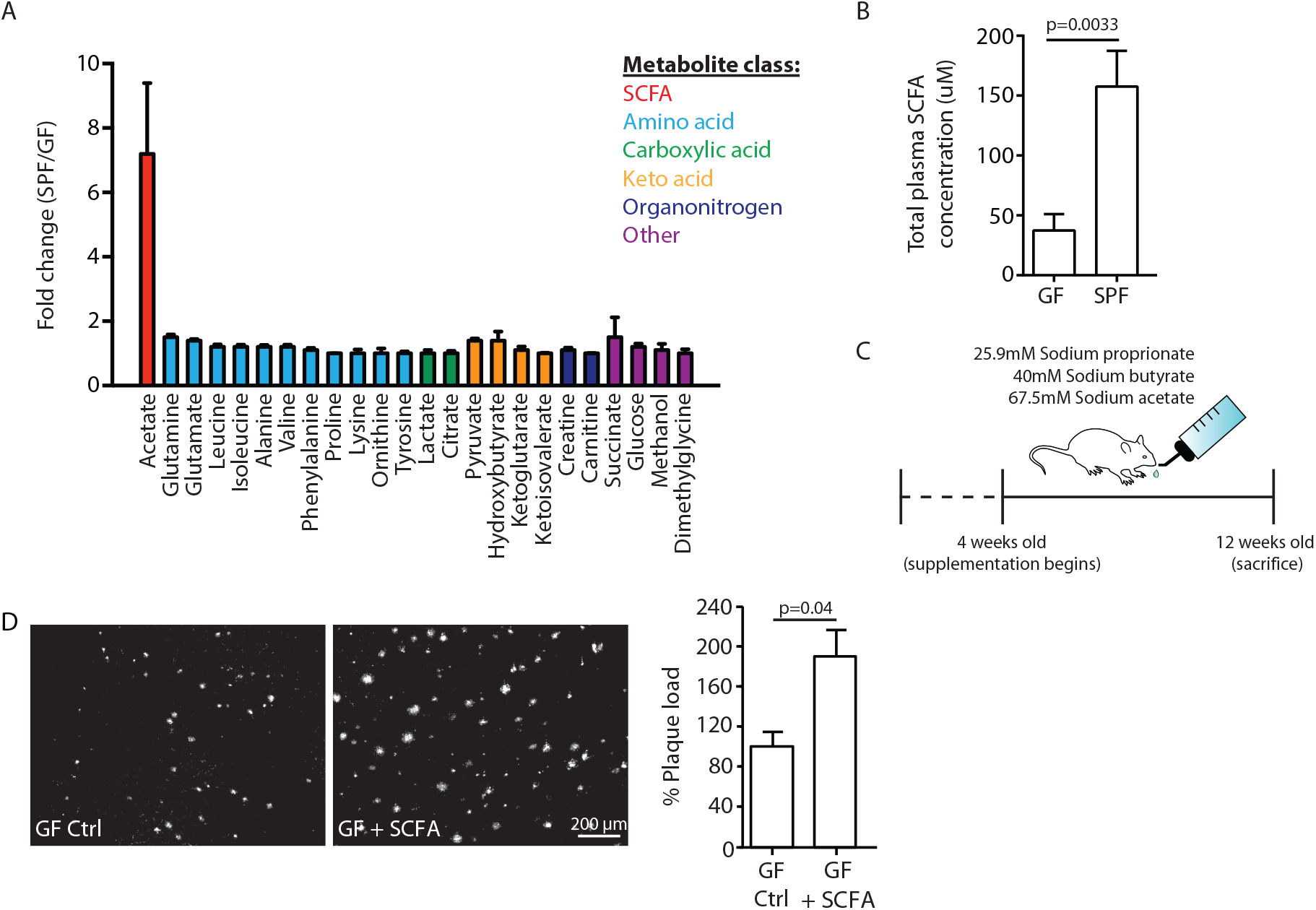
SCFA are mediators of Aβ plaque deposition. **(A)** Blood metabolome analysis showing top 15 metabolites significantly regulated between 3 months old GF and SPF APPPS1 mice. Acetate, the most abundant SCFA, was the most regulated metabolite in GF compared to SPF group. Three mice per group were analyzed. **(B)** Total plasma SCFA concentrations of acetate, butyrate and propionate showing an increase in SPF compared to GF mice (unpaired T test, n=5 per group). **(C)** Experimental plan for SCFA supplementation in GF mice. Four weeks old GF mice have been treated with SCFA in drinking water for 8 weeks. **(D)** Representative images from GF (control-treatment, ctrl) and GF supplemented with SCFA mice showing a significant increase in Aβ plaque load upon SCFA administration. Quantification of plaque area represent mean ± SEM from at least 5 mice per group (unpaired T test).

### SCFA mildly increase amyloidogenic processing

After identifying SCFA as the key and sufficient mediator of the gut microbiome’s effect on Aβ pathology, we aimed to identify the underlying mechanism. First, we explored potential direct effects of SCFA on amyloid precursor protein (APP) expression and processing by immunoblot analysis of brain tissue of GF APPPS1 mice with SCFA or control supplementation (**Fig. 3A, B**). In accordance with the histological plaque load analysis, we identified markedly increased Aβ levels in SCFA- compared to control-supplemented GF APPPS1 mice. However, levels of the full-length APP (APPFL) were comparable between the groups. Aside from a prominent and significant increase in Aβ levels, we also detected a mildly reduced ratio of the APP C-terminal fragment (CTFs) C83 (** in Fig. 3A, produced by ADAM10 cleavage) and C99 (* in Fig. 3A, produced by BACE1 cleavage) that suggested slightly increased amyloidogenic processing upon SCFA supplementation to GF APPPS1 mice. Although, this mild increase in amyloidogenic processing may add to the overall increase in Aβ, this effect alone may not be sufficient to fully explain substantially increased levels of Aβ in SCFA supplemented GF animals (**Fig. 3B**). We further analyzed protein levels of secretases responsible for APP processing (BACE1, ADAM10, and a catalytic subunit of γ-secretase PSEN1), but could not detect any alterations upon SCFA supplementation of GF APPPS1 mice (**Fig. 3A, B**). Moreover, using an in vitro γ-secretase activity assay (**Fig. 3C**), we show that addition of SCFA does not change γ-secretase activity or processivity as reflected by comparable levels of total Aβ and unaltered profiles of Aβ species (Aβ37, 38, 40 and 42/43) in this cell free assay. In contrast to a previous study (Ho et al., 2018), we could exclude a direct effect of SCFA on Aβ aggregation (**Fig. 3D**) as similar Aβ aggregation kinetics were determined in the presence or absence of SCFA in a Thioflavin T aggregation assay. Taken together, our data suggest that increased levels of Aβ triggered by SCFA *in vivo* are not mediated by major quantitative or qualitative changes in Aβ production.

**Figure 3.**
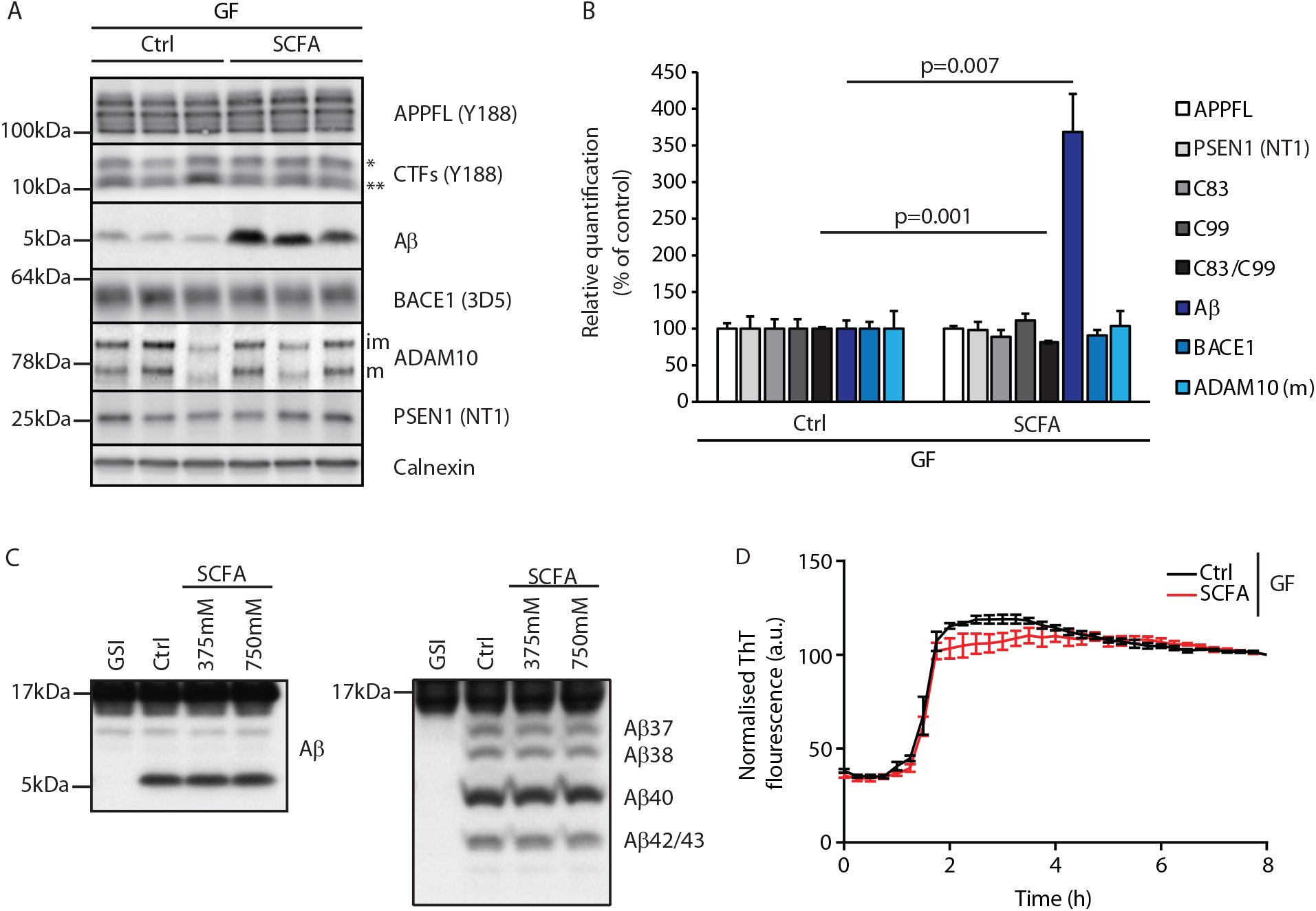
SCFA mildly increase amyloidogenic processing. **(A)** Western blot analysis and **(B)** its densitometry quantification of 3 months old brain homogenates of control (Ctrl)- and SCFA-supplemented GF APPPS1 animals. The Aβ level is significantly increased in SCFA group in comparison to Ctrl, despite unaffected APPFL levels. APP CTFs show a decreased C83 (**) to C99 (*) ratio. We could not detect alterations in protein levels of secretases involved in APP processing (ADAM10, BACE1 and γ-secretase/PSEN1). m = ADAM10 mature form; im = ADAM10 immature form. Data represent mean + SEM from 3 mice per group and p values are shown (unpaired T test). **(C)** Left panel: γ-Secretase reconstituted into lipid vesicles was incubated at 37°C together with the C99-based substrate C100-His_6_ in the presence of increasing doses of a SCFA mixture (375 and 750 μM final concentration of total SCFAs of an equimolar mixture of Na-acetate, Na-propionate and Na-butyrate) for 24h. Production of Aβ was analyzed by immunoblotting. γ-Secretase inhibitor (GSI) L-685,458 (0.4 μM) was used as a negative control. No alterations in Aβ levels were detected in the presence of SCFA. Right panel: Qualitative analysis of individual Aβ species via Tris-Bicine-Urea SDS-PAGE reveals that SCFA treatment does not alter the ratio among the different Aβ species (Aβ37-38-40-42/43) suggesting no direct effects on modulation of γ-secretase cleavage. **(D)** Aggregation kinetics of monomeric Aβ40 recorded by the increase in fluorescence of Thioflavin T incubated with either 30 mM NaCl (Ctrl) or 30 mM SCFA mixture do not show any significant difference, suggesting that SCFAs do not directly modify Aβ fibrillarization. Data points represent mean ± SD from 3 independent experiments.

### SCFA supplementation results in increased microglial activation

Given the critical role of microglia in AD and a demonstrated link between microglial function and Aβ pathology, we next focused on microglia as the potential cellular mediators of the SCFA effect on Aβ pathology (Dodiya et al., 2019). First, we observed that microglia in SCFA-supplemented GF APPPS1 mice had a significantly increased circularity index (i.e. more amoeboid shape) indicating a more activated microglial phenotype (**Fig. 4A**). We used combined single molecule fluorescent in situ hybridization (smFISH, for microglial identification by CX3CR1 expression) and immunofluorescence (for plaque identification by staining with anti-Aβ 2D8 antibody) to visualize and quantify clustering of microglia around Aβ plaques. We observed significantly more microglial cells close to plaques in SCFA- compared to control-treated APPPS1 GF mice (**Fig. 4B**). Next, we directly investigated the influence of bacterial colonization on microglial reactivity in the WT background. To this end, we injected brain homogenates from 8 months old APPPS1 mice containing abundant Aβ into the hippocampus of GF or SPF WT mice (**Fig. 4C**) and subsequently analyzed microglial abundance and activation by smFISH. We observed a significant increase in overall microglial cell counts at the peri-injection site of SPF compared to GF WT mice (**Fig. 4D**). Moreover, microglia in SPF mice expressed significantly more TREM2 mRNA puncta per microglia compared to GF WT mice (**Fig. 4E**), indicating that gut microbiome triggers microglial activation and reactivity towards an exogenous insult containing Aβ. Next, we questioned whether this SCFA-induced microglial activation might also be associated with an increased phagocytic activity of microglia, which is one of the most intensively investigated microglial functions in the context of AD. However, we did not detect any direct effect of SCFA-treatment on the phagocytic capacity of microglia using an *ex vivo* amyloid plaque clearance assay (**Fig. 4F**).

**Figure 4.**
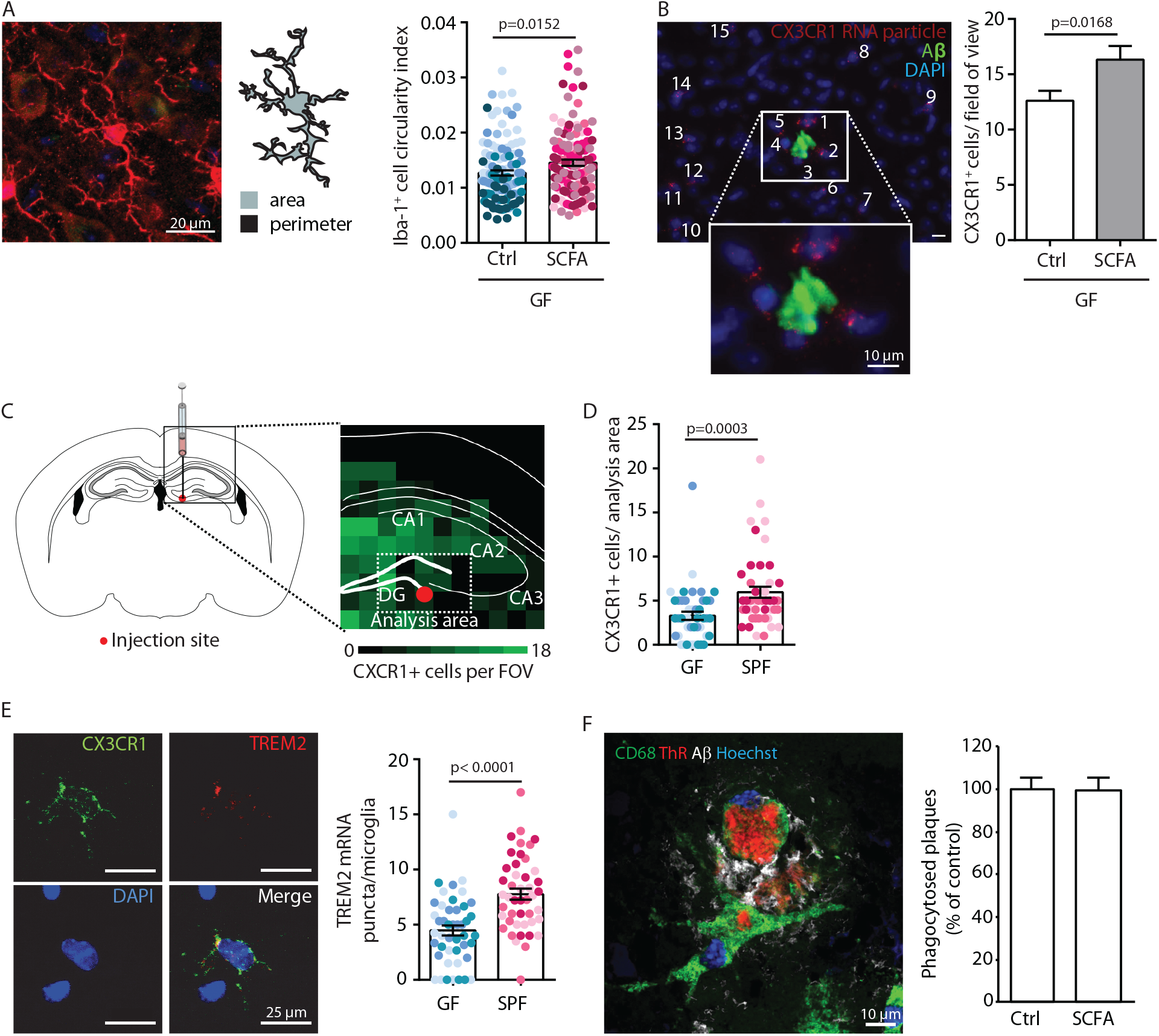
SCFA trigger microglial activation. **(A)** Morphological analysis of microglia shows an increase in the circularity index, indicating a more activated phenotype, in SCFA compared to control treated GF APPPS1 mice. Iba1 (red) has been used as microglial marker. For each group, each shade of color represents the microglia from a single mouse. **(B)** smFISH analysis of microglial cells (red, CX3CR1 mRNA particles) surrounding amyloid plaques (green, anti-Aβ clone 2D8) shows an increased number of CX3CR1-positive cells (> 4 puncta) clustering around Aβ plaques in SCFA- in comparison to control-supplemented GF APPPS1 mice. **(C)** Experimental outline for APPPS1 brain homogenate injection into a WT mouse brain showing the injection site (red dot). **(D)** smFISH analysis of CX3CR1-positive cells (>4 puncta) surrounding the APPPS1 brain homogenate injection site (analysis area relative to injection site for all brains) showing an enhanced recruitment of microglial cells in SPF versus GF WT mice. Each different shade of color represents the microglia from a single mouse. **(E)** APPPS1 brain homogenate injection induces higher microglial activation in SPF in comparison to GF WT mice as shown by the higher amount of TREM2 mRNA puncta (red) per CX3CR1-positive (green) microglia. Each different shade of color represents the microglia from a single mouse. **(F)** *Ex vivo* amyloid plaque clearance assay. Primary microglia treated with SCFA (250 μM) do not show any alteration in their phagocytic capacity towards amyloid plaques in comparison to the control treated cells (Ctrl). CD68 (green) was used to visualize microglia. Amyloid plaques were visualized using both Thiazine red (ThR, red, fibrillar Aβ) and an anti-Aβ antibody (3552, white, total Aβ). DAPI (blue) was used as nuclear dye. All data are presented as mean ± SEM from at least 3 independent experiments and at least 5 mice per group and analyzed by Mann Whitney U test (except for panel E, unpaired T test).

### Microglia-derived ApoE expression is increased upon supplementation by SCFA

To analyze the effect of SCFA on microglial function in more detail, we performed a transcriptome Nanostring analysis of brain samples from control- and SCFA-supplemented GF APPPS1 mice at 3 months of age. Corresponding to the results of the smFISH and histological analysis (**Fig. 4**), we found numerous candidate genes previously associated with microglial activation to be upregulated in SCFA-supplemented APPPS1 animals (**Fig. 5A, B**). Many of the most abundantly regulated microglial genes were associated with secretory functions (chemokines and complement factor secretion) and with pathogen recognition (e.g. Tlr7, Trem2, Tyrobp, MyD88) (**Fig. 5B**). A biological network analysis revealed upregulation of ApoE and subsequent activation of the ApoE-TREM2 pathway as a central biological pathway regulated by SCFA (**Fig. 5C**). In accordance with the transcriptomic data, we further confirmed an increase in ApoE protein expression in SCFA-compared to control-supplemented GF APPPS1 mice by immunohistochemistry (**Fig. 5D**).

**Figure 5.**
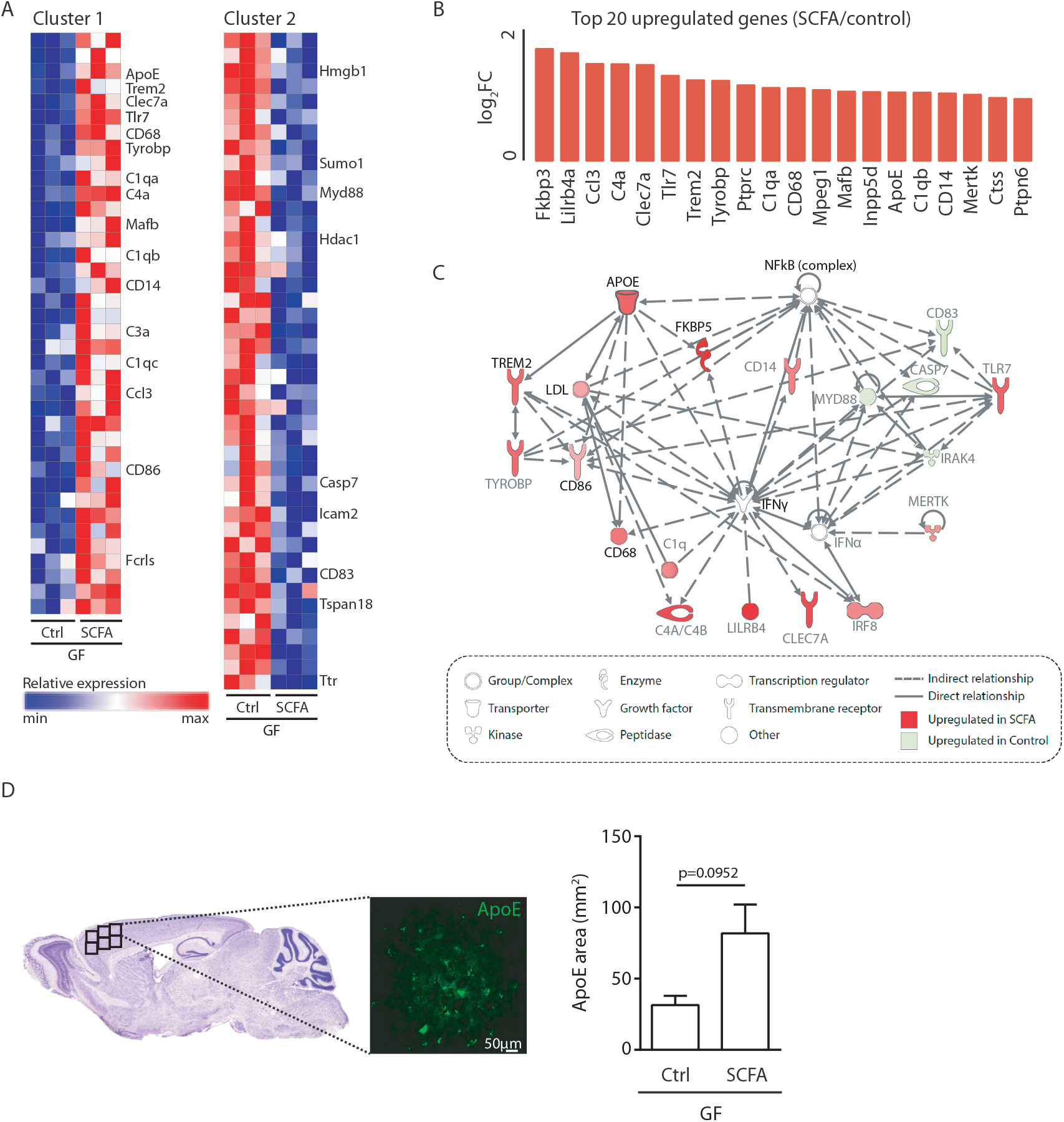
Increased ApoE expression marks microglial activation upon SCFA supplementation. **(A)** Heatmap of Nanostring transcriptomic analysis using nCounter Gene Expression mouse neuroinflammation code set from control- and SCFA-supplemented GF APPPS1 mice at 3 months of age. Row values were scaled using unit variance scaling. Genes previously associated with microglial function have been annotated on the heatmap. Three mice per group have been analysed. **(B)** Top 20 upregulated genes in SCFA versus control treated samples. Most of transcriptome hits have been previously associated with microglial activation. **(C)** Functional gene interaction network analysis using Ingenuity Pathway Analysis. Genes are colored based on fold-change values determined by RNA-Seq analysis, where red indicates an increased in SCFA- and green in control-treated animals. Network analysis revealed upregulation of the ApoE-TREM2 axis as one of the principal biological pathways upregulated by SCFA. **(D)** Representative sagittal brain section indicating location of the analyzed region of interests in the frontal cortex and representative image showing ApoE (green) distribution. Quantification of ApoE signal showed a SCFA-dependent increase of ApoE expression. Five mice per group have been analysed. Data represent mean ± SEM and p values are shown (Mann Whitney U test).

Next, we sought to investigate the cellular source of SCFA-driven ApoE expression. Previous studies have indicated astrocytes and microglia as major sources of cerebral ApoE (Holtzman et al., 2012; Shi and Holtzman, 2018). Therefore, we histologically studied expression of ApoE protein in control- and SCFA-supplemented GF APPPS1 mice and detected an increase of ApoE coverage colocalizing with microglial cells but not with GFAP-positive astrocytes (**Fig. 6A-C**). This upregulation of ApoE production by microglia was accompanied with a significant increase in ApoE-positive Aβ plaque coverage (**Fig. 6C**), indicating a potential functional role of ApoE upregulation in the SCFA-mediated increase of Aβ plaque load. Correspondingly to bacterial colonization at 5 months of age (Fig. 1D), also SCFA treatment of 3 months old APPPS1 mice resulted specifically in an increase of small plaques (4-8μm radius) (**Fig. 6D**). Taken together, these results demonstrate that SCFA supplementation results in upregulation of ApoE production specifically in microglia and this effect is associated with increased levels of ApoE in amyloid plaques and increased density of small plaques, underscoring deleterious effects of SCFA during initial stages of Aβ deposition.

**Figure 6.**
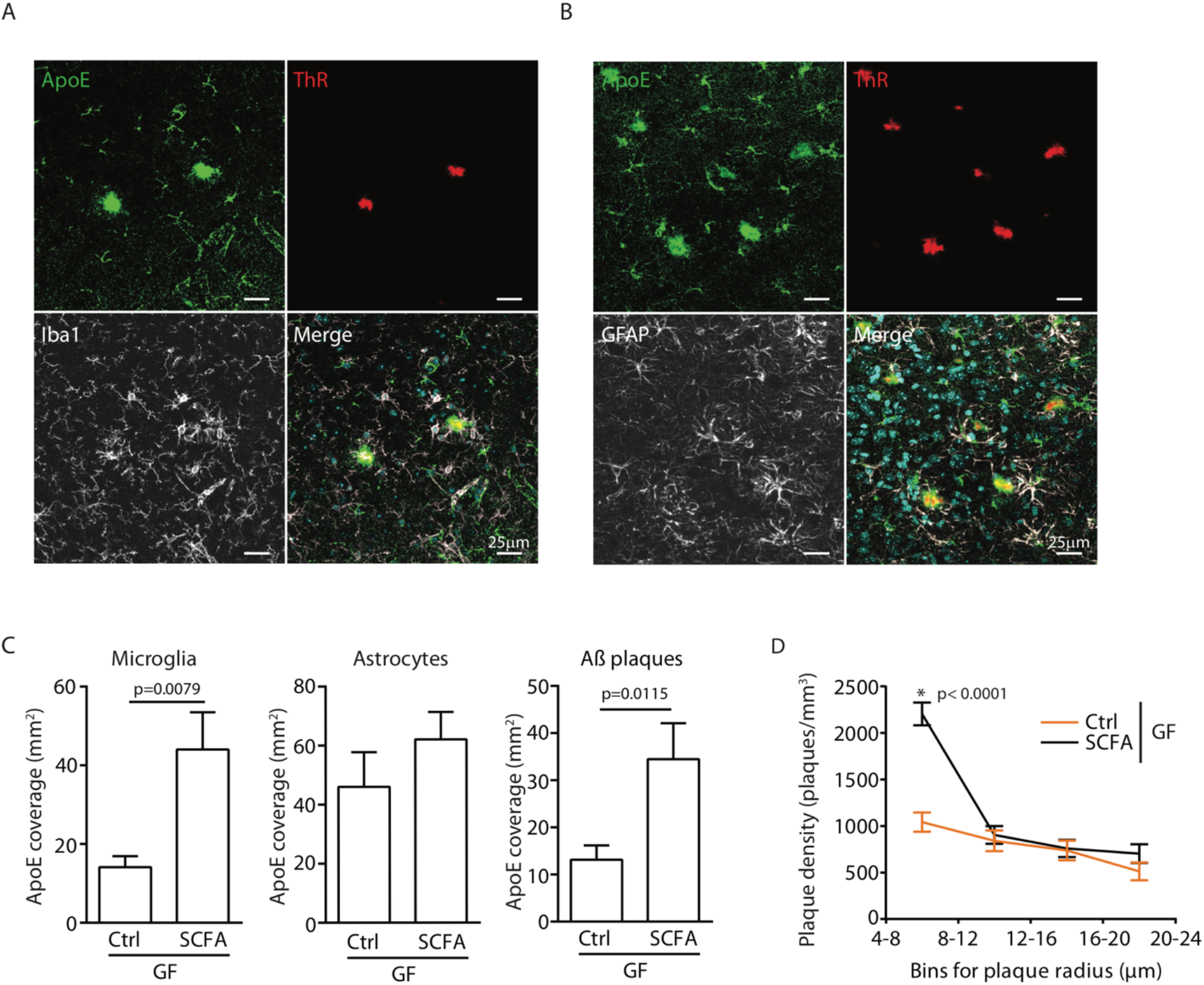
Increased ApoE expression in microglia is associated with increased levels of ApoE in amyloid plaques and increased density of small plaques. Representative images of ApoE (green) and Aβ (red, ThR) with **(A)** Iba1+ microglia (white) and **(B)** GFAP+ astrocyte (white). **(C)** Quantification of ApoE colocalization (absolute coverage area in mm^2^) with microglia, astrocytes and Aβ plaques in control- and SCFA-supplemented GF APPPS1 mice at 3 months of age. **(D)** Analysis of Methoxy-X04-stained brain sections showed a specific increase in plaques of smaller sizes (4-8 μm radius) in 3 months old SCFA- compared to Ctrl-supplemented GF APPPS1 mice. Data represent mean ± SEM from at least 5 mice per group and p values are shown (Mann Whitney U test).

## Discussion

Recent studies have demonstrated a key role of the gut microbiome in Aβ pathology in AD. However, the mechanisms by which the microbiome modulates disease progression and the molecular mediators along the gut-brain axis have remained unknown. In this study, we identified microbiota-derived SCFA as key microbial metabolites contributing to Aβ plaque deposition in the brain. GF APPPS1 mice have markedly reduced SCFA concentrations and reduced plaque load, and supplementation with SCFA is sufficient to mimic the microbiome’s effect and increase Aβ plaque burden. We identified microglia as the key cell population responsive to SCFA and increased microglial ApoE may mediate accelerated Aβ deposition during early stages of amyloidogenesis.

The gut bacteria have an intricate function in preserving the integrity of an intestinal-epithelial barrier, but exceeding this function, they have also been demonstrated to be critical for shaping the immune system, host metabolism and nutrient processing (Backhed et al., 2004; Hooper et al., 2001; Round et al., 2011). Indeed, the functions of the gut microbiome extend beyond the intestinal tract. With their rich repertoire of antigens, metabolites and direct interaction with the autonomic nervous system the microbiome potently influences remote organ function including the brain. The microbiome plays an important role in microglial maturation during development and in adulthood (Abdel-Haq et al., 2019). Previously believed to be shielded from the peripheral immune system and blood-borne mediators, it became apparent that the brain is accessible for microbial metabolites to affect cerebral immunity in health and disease (Abdel-Haq et al., 2019; Janakiraman and Krishnamoorthy, 2018).

In accordance with our observation of SCFA to be the key mediator along the gut-brain axis in AD, multiple lines of evidence have indicated that SCFA might be the key microbial metabolite group acting on brain function. Particularly, SCFA have been demonstrated to induce microglial maturation using a similar analysis paradigm by SCFA supplementation in germfree mice as used in our study (Erny et al., 2015). In a Parkinson’s disease mouse model, SCFA were sufficient to induce neuroinflammation and disease progression in absence of the gut microbiome (Sampson et al., 2016). Moreover, SCFA have been implicated in a wide range of brain disorders or physiological functions under cerebral control such as mood disorders, autism spectrum disease or energy metabolism (Cryan et al., 2019; Li et al., 2018; van de Wouw et al., 2018).

Over the past decade, microglial cells have come into the focus of experimental AD research. This has been driven by the identification of several genetic AD risk loci being associated with microglial cells (Long and Holtzman, 2019). Moreover, microglia have been demonstrated in murine AD models to actively contribute to various pathophysiological process in AD such as plaque seeding, plaque phagocytosis and neuronal dysfunction by synaptic pruning (Heneka et al., 2014; Heneka et al., 2013; Hong et al., 2016; Sarlus and Heneka, 2017). Previous proof-of-concept studies investigating a potential role of the microbiome in AD have already indicated an effect of microbial colonization or antibiotic treatment on microglial activation (Dodiya et al., 2019; Harach et al., 2017; Minter et al., 2017). Although previous studies did not reveal a mechanistic link between microbiome function to Aβ plaque pathology, they have reproducibly shown that microbiota eradication (GF mice) or impairing the microbiome’s metabolic function (antibiotic treatment) reduced microglial activation and Aβ plaque load, which is in accordance with our study. This may at first glance appear contradicting with the research hypothesis supporting microglial activation and their corresponding increased phagocytic clearance as an approach to reduce Aβ plaque load (Daria et al., 2017; Guillot-Sestier et al., 2015). However, we should bear in mind dual nature of microglial responses and their concomitant beneficial and detrimental roles in AD. Accordingly, previous studies have unequivocally demonstrated a role of microglia in promoting plaque seeding and growth, particularly at early stages of AD pathology. For example, microglia have been attributed to promote plaque seeding by release of pro-inflammatory protein complexes, so called ASC specks (Venegas et al., 2017). A recent elegant study using sustained microglia depletion starting before plaque pathogenesis revealed a critical role of microglia in formation of Aβ plaques and plaque density (Spangenberg et al., 2019). On the other hand, preventing microglial activation, such as upon loss of Trem2, increased amyloid seeding due to reduced phagocytic clearance of Aβ seeds, but at the same time reduced plaque-associated ApoE (Parhizkar et al., 2019b). Previous studies demonstrated that ApoE co-aggregates with Aβ fibrils and contributes to plaque seeding and plaque core stabilization (Liao et al., 2015; Parhizkar et al., 2019a), thus underscoring the complexity of microglia-ApoE interactions during amyloidogenesis.

ApoE is predominantly produced by astrocytes under physiological conditions, but upregulated in AD microglia (Frigerio et al., 2019). Indeed, microglial depletion in the 5xFAD mouse model resulted in a marked reduction of plaque-associated ApoE (Spangenberg et al., 2019). In the present study we identified upregulation of microglial ApoE expression. By further excluding major direct effects of SCFA on Aβ production and fibrillarization as well as on phagocytic activity of microglia, we hypothesize that increased ApoE expression may be the mechanistic link between SCFA-driven microglial activation and increased plaque load. However, final proof for this link requires investigation of SCFA effects in mice with microglia-specific ApoE depletion and awaits future studies.

SCFA use membrane-receptors as well as receptor-independent mechanisms to enter target cells and exert their functions. Receptors specifically activated by SCFA (free fatty-acid receptor-2 and −3) have been identified in several organs, including the brain (Layden et al., 2013). Yet, the SCFA can also freely enter target cells by simple diffusion or fatty acid transporters and exert their biological function intracellularly, independent of specific receptors (Lin et al., 2012; Moschen et al., 2012). Therefore, the most likely target mechanism to interfere with SCFA effects in microglia could be on the level of chromatin remodeling and regulation of target gene expression because SCFA nonspecifically inhibit class I and II histone deacetylases and induce histone acetylation (Davie, 2003; Huuskonen et al., 2004). Recent studies have shown that SCFA can modulate histone acetylation also in microglial cells and suggest that potential effects of SCFA in regulating microglial activity might be mediated via changing transcription factor binding capacities such as Nf-kB signaling (Huuskonen et al., 2004; Patnala et al., 2017). Epigenetic reprogramming of microglia is a relevant pathophysiological mechanism in microglia, and has been shown to contribute to AD pathology (Wendeln et al., 2018). However, the detailed effects of SCFA on microglial histone acetylation and the associated consequences for chromatin accessibility of inflammatory genes are currently largely unknown.

Besides direct effects on brain function including glial cell homeostasis, SCFA can also affect peripheral immune cells and thereby indirectly modulate neuroinflammatory mechanisms (Cryan et al., 2019). SCFA have been demonstrated to be potent immune modulators particularly of T cell function and their polarization into pro- and anti-inflammatory subpopulations (Smith et al., 2013). In fact, we have previously demonstrated that the effect of SCFA on post-stroke recovery in an experimental brain ischemia model was dependent on circulating T cells (Sadler et al., 2020; Sadler et al., 2017). In these previous studies, circulating T cells either mediated the effects of SCFA on the cerebral micromilieu either by their reduction in cerebral invasion or by polarization of the secreted cytokine profile. However, in contrast to AD, stroke induces a pronounced neuroinflammatory response to the acute tissue injury with the invasion of large numbers of circulating T cells, which have been demonstrated to contribute substantially to stroke pathology (Cramer et al., 2019; Liesz et al., 2009). Therefore, while the peripheral polarization of T cells and their consecutive impact on local neuroinflammation is conceivable after stroke, this pathway seems less likely in AD with only very limited invasion of circulating immune cells into the brain, particularly in the early stages of AD pathology.

Interestingly, the biological effect of SCFA on microglia seems to be largely dependent on the specific disease condition. While SCFA in neurodegenerative conditions, as also in our study, have been associated with microglial reactivity and activation, their function in primary autoimmune and acute brain disorders has mainly been described to be anti-inflammatory. For example, we have been previously shown that SCFA treatment promotes anti-inflammatory mechanisms and improve neuronal function in ischemic stroke and similar findings have also been reported in experimental autoimmune encephalitis (Haghikia et al., 2015; Sadler et al., 2020). The detailed cause of this divergent functions of SCFA in different disease conditions is currently still unknown and requires further exploration for the potential use of personalized treatment approaches. Yet, in light of our results in the AD compared to the stroke model, it seems likely that the different effects of SCFA treatment on disease outcome could be due to engaging different mechanistic routes, such as peripheral lymphocyte polarization in EAE and stroke versus local effects on microglia in AD.

Our study suggests that SCFA-regulated pathways might be promising drug targets in the peripheral circulation for early-stage AD to prevent microglial activation, ApoE production and the development of amyloid pathology. However, the therapeutic targeting and neutralization of SCFA, e.g. by specific SCFA-scavengers, in order to chronically reduce circulating SCFA blood concentrations is currently not established. Also attempting to reduce SCFA concentrations by reduction of nutritional fiber intake is not a feasible therapeutic approach. Reduced fiber intake correlates with increased risk of metabolic syndrome and cardiovascular events such as myocardial infarction and stroke. Furthermore, dietary fiber restriction will most likely not be efficient to affect microglial activation in early-stage AD because a near-complete reduction of blood SCFA concentrations, as seen in GF mice, will not be achieved by dietary intervention.

In conclusion, our study identifies SCFA as the key molecular mediators along the gut-brain axis in AD. Identification of this novel pathway will open up new avenues for therapeutic targeting of the microbiome-SCFA-microglia axis to reduce the inflammatory impact on AD development.

## Materials and Methods

### Animal experiments

All animal experiments were performed under the institutional guidelines for the use of animals for research and were approved by the governmental ethics committee of Upper Bavaria (Regierungspraesidium Oberbayern). SPF B6.Cg-Tg (Thy1-APPSw,Thy1-PSEN1*L166P)21Jckr (APPPS1) (Radde et al., 2006a) mice were originally donated by Matthias Jucker (Tübingen, Germany) to Christian Haass (Munich, Germany) and bred for this project at the core animal facilities of the Center for Stroke and Dementia Research in Munich. Mice were kept at 12h dark-light cycle with ad libitum access to food and water.

### GF mouse generation and handling

APPPS1 mice were rederived to GF status in the Clean Mouse Facility, University of Bern, Switzerland as previously reported (Harach et al., 2017) and housed in flexible-film isolators. All mouse handling and cage changes were performed under sterile conditions. GF and SPF mice all received the same autoclaved chow and sterile water. For SCFA treatment, mice were given a sterile-filtered solution containing 25.9 mM sodium propionate, 40 mM sodium butyrate and 67.5 mM sodium acetate in sterile water ad libitum from 4 weeks of age. The SCFA water solution was renewed every three days. For surgical interventions (stereotactic injection, see below) in GF mice, the whole surgical procedure and post-surgical care was performed in a microbiological safety cabinet as previously described in detail (Singh et al., 2018). Animals were regularly checked for germfree status by aerobic and anaerobic cultures, cecal DNA fluorescence stain and by 16s rRNA PCR of fecal pellets. GF status of all animals used in this study (control and SCFA-supplemented) has been confirmed after sacrificing the mice.

### Histological analysis of Aβ plaque load and density

Mice were transcardially perfused with PBS followed by overnight post fixation with 4% PFA solution. Free floating 30 μm sagittal brain sections have been permeabilized and blocked for 1h in PBS/0.5% Triton x-100/5% normal goat serum (NGS). Next, samples have been incubated overnight at 4°C with primary antibody anti β-amyloid (Aβ) (clone 6E10, 1:500, BioLegend) diluted in blocking buffer and stained with the corresponding goat secondary antibody. Immunostainings have been performed on 6 brain sections/animal collected every 300 μm starting from the interhemispheric fissure. Three 10x images/section (front, middle and rear cortex) have been acquired and the plaque load (6E10 coverage area) has been determined using the particle analysis tool in ImageJ software (NIH) and normalized on total tissue area. Analysis has been performed at least on 5 mice per group/time point. Histological analysis has been performed by an investigator (A.-V. C) blinded to treatment groups. For analysis of plaque density, 30 μm sections were stained with Methoxy-X04 and confocal images were collected in Z-stacks. 8 ROI were selected across the cortex and acquired across the different cortical layers. Image data analysis was performed with custom-written MATLAB script as previously reported (Peters et al., 2018). In brief, local background correction was applied to diminish intensity variations among different stacks and to account for the intensity decline in the axial dimension due to absorption and scattering of photons. For this purpose, the voxel intensity was normalized in each Z-layer to the 70th percentile of Methoxy-X04 fluorescence intensity. Subsequently, amyloid plaques were defined by applying the 90th percentile in the Methoxy-X04 fluorescence data stacks. The radius of each individual plaque was calculated from the Z-plane with the largest area extension in XY (radius=√(area/π)), assuming spherical shape of plaques (Hefendehl et al., 2011). All plaques that contacted the image border were excluded from the analysis. The cut-off size was set to a minimal plaque radius of 2 μm.

### Barnes maze test for memory deficits

A modified version of Attar A et al (Attar et al., 2013) was used to perform the Barnes maze test. The elevated 20-hole apparatus (diameter: 100 cm, hole diameter: 10cm) had a target box which was placed under the maze. The protocol includes three phases of interaction of mice with the maze (1) habituation, (2) 2-day training and 48 h later (3) probe. Before each day of training or probe mice were placed 30 min prior procedure in the testing room for acclimatization. On day 1 mice were habituated to the maze. Therefore, mice were placed in the center of the maze in a 2l glass beaker. After 1 min of acclimatization, mice were guided slowly by moving the glass beaker towards the target hole. This was done three consecutive times per mouse. On day 2 mice were placed in center and given the possibility to freely explore and find the target hole. If mice did not reach the target hole within 3 min the glass beaker was used to slowly guide them to the target. This was done in three consecutive trials. On day 3 mice were placed under the beaker and 10 s after placement the beaker was removed, and mice were allowed to explore freely and find the target hole. Again, three consecutive trials were performed with every mouse. This procedure was repeated for the actual probe 48 h later. Acquisition and zone-dependent analysis were performed with Ethovision XT (Noldus). Data acquisition and analysis was performed by an investigator (S.R.) blinded to treatment groups.

### Metabolomics analysis by NMR

#### Sample preparation

A 75 mM disodium phosphate buffer solution in H2O/D2O (80/20) with a pH of 7.4 containing 6.15 mM NaN_3_ and 4.64 mM sodium 3-[trimethylsilyl] d4-propionate (Cambridge Isotope Laboratories) was prepared. After thawing of the frozen mouse plasma samples, 125 μl of buffer and 125 μl of plasma were manually transferred into a Ritter 96 deep well plate. The samples were mixed by manually aspirating and dispensing 3 times. Using a modified Gilson 215 liquid handler, 195 μl of each sample were transferred into 3 mm NMR SampleJet tubes. The tubes were closed by inserting POM balls into the caps and subsequently transferred to the Sample Jet autosampler were they were kept at 6 °C while queued for acquisition.

#### NMR experiments and processing

All proton nuclear magnetic resonance (^1^H-NMR) experiments were acquired on a 600 MHz Bruker Advance II spectrometer (Bruker BioSpin, Karlsruhe, Germany) equipped with a 5 mm triple resonance inverse (TCI) cryogenic probe head with Z-gradient system and automatic tuning and matching. A standard 3mm sample of 99.8% methanol-d4 (Bruker Biospin) was used for temperature calibration (Findeisen et al., 2007) before the measurements. All experiments were recorded at 310 K.

The durations of the π/2 pulses were automatically calibrated for each individual sample using a homonuclear-gated nutation experiment (Wu and Otting, 2005) on the locked and shimmed samples after automatic tuning and matching of the probe head. For water suppression, pre-saturation of the water resonance with an effective field of γB_1_ = 25 Hz was applied during the relaxation delay.

T2-filtered 1H NMR spectra were acquired with the standard 1D Carr-Purcell-Meiboom-Gill (CPMG) pulse sequence was used with a relaxation delay of 4 seconds. A pulse train of 128 refocusing pulses with individual spin echo delays of 0.6 ms was applied resulting in a total T2 filtering delay of 78 ms. after applying 4 dummy scans, a total of 73,728 data points covering a spectral width of 12,019 Hz were collected using 16 scans. J-resolved spectra (JRES) were recorded with a relaxation delay of 2 seconds and a total of 1 scan for each increment in the indirect dimension after 8 dummy scans. A data matrix of 40 × 12,288 data points was collected covering a sweep width of 78 × 10,000 Hz. The time-domain data was automatically processed and the metabolites quantified using a version the KIMBLE metabolomics workflow adapted for plasma NMR measurements (Verhoeven et al., 2018).

### Sample preparation for biochemical analyses

Mouse brains were isolated and immediately snap frozen in liquid nitrogen. Frozen brains were then mechanically pulverized for further applications and stored at −80 °C. Aliquots of brain powder were lyzed for 20 min in lysis buffer (150 mM NaCl, 50 mM Tris pH 7.5, 1% Triton X-100) supplemented with protease and phosphatase inhibitor cocktail (Roche) on ice. Samples were then centrifuged at 17,000 g for 30 min at 4 °C. Supernatants were collected (soluble fraction) and stored at −80 °C.

### In vitro amyloid aggregation assay

Synthetic Aβ40 peptide (rPeptide) was solubilized (1 mg/ml) in 50mM NH4OH (pH >11). Upon 15min incubation at RT and subsequent water bath sonication (5 min), fractions of 100 μg Aβ40 were lyophilized and stored at −20°C. 10 μg Aβ40 was suspended in 20 mM NaPi with 0.2 mM EDTA buffer (pH 8.0). After water bath sonication, the Aβ40 solution was filtered through an Anatop 0.02 μm filter and stored on ice. Final Aβ40 concentration was assessed from the UV spectrum ([Aβ40] = (A275-A340)/1280 mol/l). Circular Dichroism measurements (Jasco 810 Spectropolarimeter) confirmed the random coil conformation of the Aβ peptide. For the rtQuic aggregation experiments, Aβ40 was diluted to 20 μM final concentration in 20 mM NaPi, 0.2 mM EDTA buffer (pH 8.0), containing 100 μM Thioflavin T. 50 μl Aβ40 solution was either mixed with 50 μl SCFA mixture (i.e. 20 mM Na-acetate + 20 mM Na-propionate + 20 mM Na-butyrate, 100 μM Thioflavin T) or mixed with 50 μl saline solution (60 mM NaCl, 100 μM Thioflavin T), to compare aggregation of Aβ40 (final 1 μM) in 20 M NaPi, 0.2 M EDTA, pH 8.0, either enriched with 30 mM SCFA mixture or with 30 mM NaCl. Experiments were performed in triplicates. Aβ40 solution aliquots were incubated at 37 °C in 96-well plates under constant double-orbital shaking in a FLUOStar Omega plate reader (BMG Labtech). The formation of amyloid fibrils was monitored by the Thioflavin T fluorescence. Data were assessed every 15 min (λex = 440 nm, (λem = 480 nm). In control experiments (blanks), 100 μM Thioflavin T was diluted in the 20 mM NaPi, 0.2 mM EDTA, pH 8.0 buffer alone, or enriched with the SCFA mixture (30 mM) or saline solution (30 mM NaCl).

### Immunoblot analysis of mouse brain

For western blot analysis, the soluble fraction from 3 months old APPPS1 mice have been quantified using Bradford assay (Biorad) according to manufacturer’s protocol. At least 10 μg per sample have been loaded either on a bis-tris acrylamide (APP, NT, ADAM10 and BACE1) or a Novex 10-20% Tris-Tricine gel (Aβ, C83 and C99) followed by blotting on nitrocellulose membrane (Millipore) using the following antibodies: anti-APP-Cterm (Y188, 1:1000, Abcam), anti-Aβ (3552, 1:1000, (Yamasaki et al., 2006)), anti-Presenilin 1 (N terminus) (NT, 1:1000, BioLegends), anti-ADAM10 N-terminal (1:1000; R&D Systems) and anti-BACE1 (1:1000, Epitomics). Blots have been developed using horseradish peroxidase-conjugated secondary antibodies (Promega) and the ECL chemiluminescence system (Amersham). An antibody against calnexin (1:1000, Stressgen) has been used as loading control. At least 3 mice per group was analyzed.

Densitometry analysis has been performed using gel analyzer tool on ImageJ (NIH).

### γ-Secretase activity assay

γ-Secretase cleavage assays using C100-His6 as substrate were carried out using purified γ-secretase reconstituted into small unilamellar vesicles (SUV) composed of POPC essentially as described (Winkler et al., 2012) except that the reconstitution was performed in 0,5x PBS and 30 mM DTT. The SCFA mixture (Na-acetate, Na-propionate and Na-butyrate dissolved in water in equimolar ratios) was added to the assay samples at the indicated final concentrations from stock solutions. Following separation of the assay samples by SDS-PAGE on Tris-Tricine or Tris-Bicine urea gels (for total Aβ or Aβ species, respectively) (Winkler et al., 2012), Aβ was detected by immunoblotting using antibody 2D8 (Shirotani et al., 2007).

### Immunohistological microglia analysis

Free floating 30 μm sagittal brain sections were treated for antigen retrieval using sodium citrate buffer and subsequently stained using primary antibodies for Iba1 (1:250, rabbit, Wako) and anti-6E10 (1:1000, Biolegend), and the corresponding secondary antibodies and nuclear counterstain using DAPI. Images were acquired in the frontal cortex using a Zeiss confocal microscope with 40x magnification. Morphological analysis of microglial cells was performed using FIJI software. Each single microglial cell was identified in the Z-stack and a maximum intensity projection (MIP) was created for each individual cell from the image layers covering the individual microglial cell and the resulting image was binarized by thresholding. The area and the perimeter of the cell shape were measured and the circularity index (CI) was calculated (CI = 4π[area]/[perimeter]2) for each cell.

For analysis of ApoE expression in microglia and astrocytes, sections were stained with anti-P2Y12 (1:100, Anaspec #AS-55043A) or anti-GFAP (1:200, DakoCytomation Z 0334), mouse anti-6E10 antibody (1:200, abcam #16669) and biotin anti-ApoE antibody (1:50, abcam #16669). Slides were then labelled with the secondary antibodies and counterstained with DAPI. Samples were imaged using an epifluorescence microscope (40x magnification) and co-localization of Aß, ApoE, microglia was analyzed using FIJI software.

### Stereotaxic hippocampal injections

All animal handling and surgical procedures for GF as well as the SPF mice were performed under sterile conditions in a microbiological safety cabinet. Animals were anaesthetized, placed in a stereotactic device and a midline incision was performed to expose the skull. The skull was thinned above the injection point (coordinates: lateral 1mm, caudal 2.3mm from bregma) and 2.5 μl mouse brain homogenate with high plaque burden from 8 months old APPPS1 mice or saline were microinjected (Hamilton syringe at 1μl/min; 2.5mm depth from the brain surface). The needle was retained for additional two minutes before it was slowly withdrawn. 24 h after injection, animals were sacrificed, and brain samples processed for immunofluorescence and fluorescence in situ hybridization.

### Single-molecule fluorescence in situ hybridization

Single-molecule fluorescence in situ hybridization (smFISH) was performed using the RNAscope Multiplex Fluorescent Reagent Kit v2 (Advanced Cell Diagnostics) by the manufacturer’s protocols. Briefly, free floating 30 μm sagittal brain sections were first dried, washed and then incubated in RNAscope Hydrogen Peroxide. Antigen retrieval and protease treatment was performed as per protocol. Sections were then incubated with the probe mix (C2-TREM2 and C1-CX3XR1) for 2 h at 40 °C and then immediately washed with wash buffer. Next, sections were incubated with RNAscope Multiplex FL v2 AMP1, AMP2 and AMP3, and then probes were counter stained with TSA Plus Cy3 for C1-CX3CR1 and TSA Plus Cy5 for C2-TREM2. After washing with PBS and blocking buffer, plaques were identified using primary rat anti-2D8 antibody (1:300, Abcam #16669) and counter stained with secondary antibody AF488 goat anti-rat, (1:200, Invitrogen #A11006). Finally, sections were stained with DAPI. smFISH-stained RNA molecules were counted only within the DAPI staining of the cell; a cell was considered CX3CR1-positive when more than four CX3CR1 puncta were present. A cell was quantified as TREM2 positive when TREM2 smFISH molecules were observed in the CX3XR1-positive cells.

### Ex vivo Aβ plaque clearance and phagocytosis assay

We investigated microglia phagocytosis by an assay that we have previously established (Bard et al., 2000). Briefly, 10 μm brain sections from APPPS1 mice (Radde et al., 2006b) were incubated on coverslips with anti-Aβ antibodies (6E10, 5μg/ml) to stimulate microglia recruitment. Primary microglia were isolated from P7 WT mouse pups using the Neural Tissue Dissociation Kit, CD11b microbeads and a column MACS separation system (Miltenyi Biotec). Purified microglia were resuspended in DMEM F12 with 10% fetal bovine serum and 1% PenStrep (Gibco). Then, isolated microglia were incubated for 5d at a density of 3×10^5^ cells/coverslip in culturing medium containing 250 μM mixed SCFA (83.3 μM acetate, butyrate and propionate, each) or control 250 μM NaCl. Next, sections were fixed and permeabilized and stained with primary antibodies against CD68 (1:500, AbDserotec) and Aβ (3552, 1:500, (Yamasaki et al., 2006)) and corresponding secondary antibodies as well as the nuclear dye Hoechst and fibrillar Aβ dye Thiazine red (ThR 2 μM, Sigma Aldrich). Full sections were analyzed by tile scan mode on a Leica SP5 II confocal microscope. To evaluate microglial phagocytic capacity, difference in plaque coverage (ThR signal area) was quantified between brain sections incubated with microglia and a consecutive section where no microglia have been plated using ImageJ software (NIH). Each experimental group has been tested in at least 3 independent experiments with 2 technical replicates each.

### Nanostring analysis

Brain samples were lysed in Qiazol Lysis Reagent and total RNA was extracted using the MaXtract High Density kit with further purification using the RNeasy Mini Kit (all Qiagen). 70 ng of total RNA per sample was then hybridized with reporter and capture probes for nCounter Gene Expression code sets (Mouse Neuroinflammation codeset) according to the manufacturer’s instructions (NanoString Technologies). Samples were injected into NanoString cartridge and measurement run was performed according to nCounter SPRINT protocol. Background (negative control) was quantified by code set intrinsic molecular color-coded barcodes lacking the RNA linkage. As positive control code set intrinsic control RNAs were used at increasing concentrations. Data were analyzed using the nSolver software 4.0. In the standard data analysis procedure, all genes counts were normalized to the positive control values and the values for the standard reference genes. Clustering and heatmaps were performed using the ClustVis package on normalized expression values derived from the nSolver report (Metsalu and Vilo, 2015). For clustering, the z-scores were calculated using the mean expression of biological replicates per condition and then clustered using K-means. Data were further analyzed using the Ingenuity software (Ingenuity Systems). Differentially expressed genes (with corresponding fold changes and p values) were used for generating a merged biological network as previously described (Butovsky et al., 2014).

### Statistical analysis

All analyses, unless otherwise specified, have been performed using GraphPad Prism software (GraphPad, version 6.0). Sample size was chosen based on comparable experiments from previous experiments. For experimental design details on sample size, please refer to manuscript results and figure legends. A p value of <0.05 was regarded as statistically significant.

## Acknowledgments

The authors would like to thank Kerstin Thuß-Silczak for excellent technical support and Anna Daria for the help in genotyping. The authors are grateful to Mathias Jucker (Hertie-Institute for Clinical Brain Research, University of Tübingen, Germany) for providing the APPPS1 mice and David Holtzman (Washington University School of Medicine, St Louis, Missouri, USA) for providing the ApoE antibody. This work was funded by the Vascular Dementia Research Foundation, the European Research Council (ERC-StG 802305), an EU H2020-NMP-2015-686271 grant, the Munich Cluster for Systems Neurology (EXC 2145 SyNergy – ID 390857198), the German Research Foundation (DFG, STE847/6-1 and HA1737/16-1), the Helmholtz Society (Zukunftsthema “Immunology and Inflammation”), the NCL Foundation and the Alzheimer Forschung Initiative e.V.

## Author contributions

AVC and RKS performed the majority of experiments and contributed to writing the manuscript; AVC, RKS, GL, VS, SR, SH, LSM, AV, FP, SP, FK, MG performed experiments and analyzed data; MG, AM, KM, CB, MD and CH provided essential material and equipment for this study and provided critical input to manuscript writing; HS, MAG, CH, ST and AL supervised experiments and analyzed data; ST and AL initiated and coordinated the study and wrote the manuscript.

